# *Mycobacterium tuberculosis* suppresses protective Th17 responses during infection

**DOI:** 10.1101/2025.05.08.652811

**Authors:** Alex Zilinskas, Amir Balakhmet, Douglas Fox, Heyuan Michael Ni, Carolina Agudelo, Helia Samani, Sarah A. Stanley

## Abstract

*Mycobacterium tuberculosis* (Mtb) causes more deaths annually than any other pathogen, yet an effective vaccine remains elusive. IFN-γ–producing Th1 CD4+ T cells are necessary but insufficient for protection against infection. In humans, the development of IL-17A producing Th17 T cells correlates with protection, however not all individuals develop a Th17 response. In mice, experimental vaccines can elicit protective Th17 cells, yet Th17s are rare in primary infection. Why Mtb fails to consistently elicit Th17s is unknown. Here, we identify factors suppressing Th17 responses during primary infection. We demonstrate that the lack of Th17 induction is independent of route and duration. Next, using *Tbet* deficient mice, we show that Mtb drives a Th1 response that is only partially protective and limits Th17 cell production in an IFN-γ independent manner. We further reveal that the ESX-1 type VII secretion system and lipid PDIM suppresses Th17 responses. Infection with ESX-1 or PDIM mutants results in significantly increased Th17 T cells and IL-17A cytokine in the lungs, and infection of IL-17A deficient animals partially restores virulence of ESX-1 and PDIM mutants. Although the ESX-1 secretion system and lipid PDIM elicits type I IFN, which can suppress Th17 differentiation, we find that suppression of Th17 is independent of type I IFN. Instead, ESX-1 and PDIM suppresses production of IL-23, a cytokine that promotes Th17 differentiation, in dendritic cells found in mediastinal lymph nodes during Mtb infection. These findings define a new function of the ESX-1 secretion system and PDIM in Mtb virulence, a long-standing question in tuberculosis research.

## Introduction

*Mycobacterium tuberculosis* (Mtb), the causative agent of tuberculosis (TB), remains one of the leading infectious disease killers worldwide, with 10.8 million new cases and 1.25 million deaths reported in 2021 (1). The current vaccine, *Mycobacterium bovis bacillus Calmette-Guérin* (BCG), protects against childhood TB, but has highly variable efficacy, wanes over time, and provides minimal protection against adult pulmonary TB (1–3). Although there are 17 vaccine candidates currently in clinical trials, achieving robust vaccine elicited immunity remains a major challenge (4).

Interferon-γ (IFN-γ) produced by type 1 T helper (Th1) cells is essential for control of Mtb infection both in animal models and in humans (5, 6). However, the MVA85A vaccine, despite inducing robust Th1 responses as intended, failed to provide protection against Mtb infection (7). Patients with active TB can have high levels of IFN-γ in the lungs despite being unable to control infection (8). Thus, Th1 cells are necessary but not sufficient for controlling Mtb infection suggesting that other factors are important for effective immunity. Without a comprehensive understanding of which types and specific subsets of T helper cells are essential to limit Mtb infection, developing an effective Mtb vaccine with consistent high efficacy will remain elusive.

Type 17 T helper (Th17) cells are known for their association with mucosal sites, including the lungs, where they have been shown to provide protection against several bacterial and fungal respiratory pathogens (9). This protection relies on IL-17A production by Th17 cells (10). Many factors determine whether Th17 cells are elicited by infection. Antigen-presenting cells (APCs) drive the polarization of naïve T cells into Th17 cells by producing cytokines including IL-1β, IL-6, TGF-β, and IL-23. Activation of the RORγT transcription factor is essential for Th17 polarization (11). There is also evidence that the route of infection influences whether a Th17 response is elicited as infection with *Listeria monocytogenes* via the intranasal, but not intravenous route is required to observe a Th17 response to infection (12).

Th17 differentiation is opposed by Th1 differentiation via multiple mechanisms. Tbet, the transcription factor essential for commitment to the Th1 lineage, has been shown to prevent Runx1 activation, thereby preventing Runx1-mediated transactivation of RORγT and Th17 differentiation (11). Additionally, IFN-γ, the signature cytokine of Th1s, inhibits Th17 differentiation and function via activation of STAT1, independent of Tbet (13). IFN-γ inhibits Th17 differentiation and function via Tbet-dependent and Tbet-independent mechanisms (13). Finally, production of type I IFN also can suppress Th17 differentiation (14). Thus, several hallmarks of Mtb infection, namely strong Th1 responses, IFN-γ production, and type I interferon production could limit differentiation of Th17s.

Increasing evidence that Th17 cells correlate with protection against Mtb infection both in the non-human primate model (15) and in humans (16–18) suggest that Th17 cells are an important component of protective immunity to Mtb. In mice, Th17 cells are rarely observed during primary Mtb infection, and the conditions under which Th17 cells are induced during infection remain unclear (19). However, experimental TB vaccines, particularly when administered via a mucosal route, elicit Th17 cells that can confer protection against subsequent challenge in mice (20–22), demonstrating that Th17 cells can be protective against Mtb infection. Why Mtb infection does not robustly elicit Th17 cells during primary infection, despite causing infection at mucosal surfaces via airway infection, is unclear.

The ESX-1 type VII secretion system is a fundamental determinant of Mtb virulence (23). The ESX-1 secretion system has been demonstrated to subvert and/or manipulate immune responses, including by perforating phagosomal membranes, inducing detrimental host type I interferon, and inducing autophagy (24–26). As much of the work examining ESX-1 function has been conducted in vitro, we lack a clear understanding of all the factors that attenuate ESX-1 mutants during in vivo infection. Interestingly, the Mtb lipid phthiocerol dimycocerosate (PDIM) is also required for virulence and has been linked to ESX-1 function through induction of type I IFN, which can suppress Th17 responses (27, 28). Furthermore, the ESX-1 secretion system was shown to suppress production of IL-12 p40 (23), a subunit shared with IL-23, which promotes Th17 differentiation. These data suggest that Mtb virulence factors, including ESX-1 and PDIM, may suppress the emergence of a protective Th17 response during Mtb infection.

Here we show that the induction of Th17 cells during Mtb infection is suppressed by multiple mechanisms. First, we show that the absence of a Th17 response during primary infection is not dependent on duration or route of infection. Instead, we show that mice lacking Tbet (*Tbx21*^−/−^) produce Th17 cells after Mtb infection, and that these cells provide IL-17A-dependent protection against early Mtb infection. This suppression of Th17 production by Th1 cells is independent of IFN-γ. Furthermore, we observe a strong Th17 response during infection with ΔESX-1 or PDIM-lacking Mtb in mice, suggesting that these virulence factors function to suppress Th17 differentiation. Surprisingly, the ability of ESX-1/PDIM to suppress Th17 differentiation is not related to their enhanced induction of type I IFN. Instead, we show that during infection with ΔESX-1 or PDIM-lacking Mtb strains type 2 conventional dendritic cells (CD11b^+^ DCs, cDC2s), which are known to support Th17 differentiation, produce higher levels of IL-23 secretion in draining lymph nodes of Mtb infected mice (29). These results demonstrate that Mtb utilizes the ESX-1 secretion system and PDIM to bias host T helper response towards Th1s and away from Th17s.

## Results

### Th17 induction during *M. tuberculosis* infection does not depend on duration or route of infection

We first tested whether the duration or route of infection has an impact on the generation of Th17 cells during Mtb infection. To test for induction of Th17 cells, we infected wild type mice with ∼100 CFU of WT *M. tuberculosis* via the aerosol route, a standard dose for the mouse model (30). As previously published by multiple groups, we do not observe Th17 cells during the first month after Mtb infection of mice (19–22). At 4 weeks post infection, we observed the expected plateau of CFU in the lungs of infected mice indicating the onset of T cell-based control of infection (Figure 1A). With stimulation using peptide pools from model antigens ESAT-6 and Antigen 85B (Ag85B) (31–33), we observed only IFN-γ^+^ CD4^+^ (Th1) T cells (Figure 1B) and no IL-17A^+^ CD4^+^ (Th17) T cells at this or earlier timepoints (Figure 1C). Th17 cells have been observed in humans, at times presumably long after initial infection (16–18). We therefore tested whether Th17 cells arise at late timepoints after infection in mice. Mice continued to have a Th1 response up to day 170 post infection, with no increase in Th17 levels observed (Figures 1D-E). It was previously reported that infection with the virulent HN878 “Beijing” strain of the L2 lineage resulted in induction of Th17 cells (19). However, we did not observe a significant increase in IL-17A producing T cells in murine lungs when we infected with a Beijing strain at 3 weeks post infection (Supplementary Figure S1).

**Figure 1.**
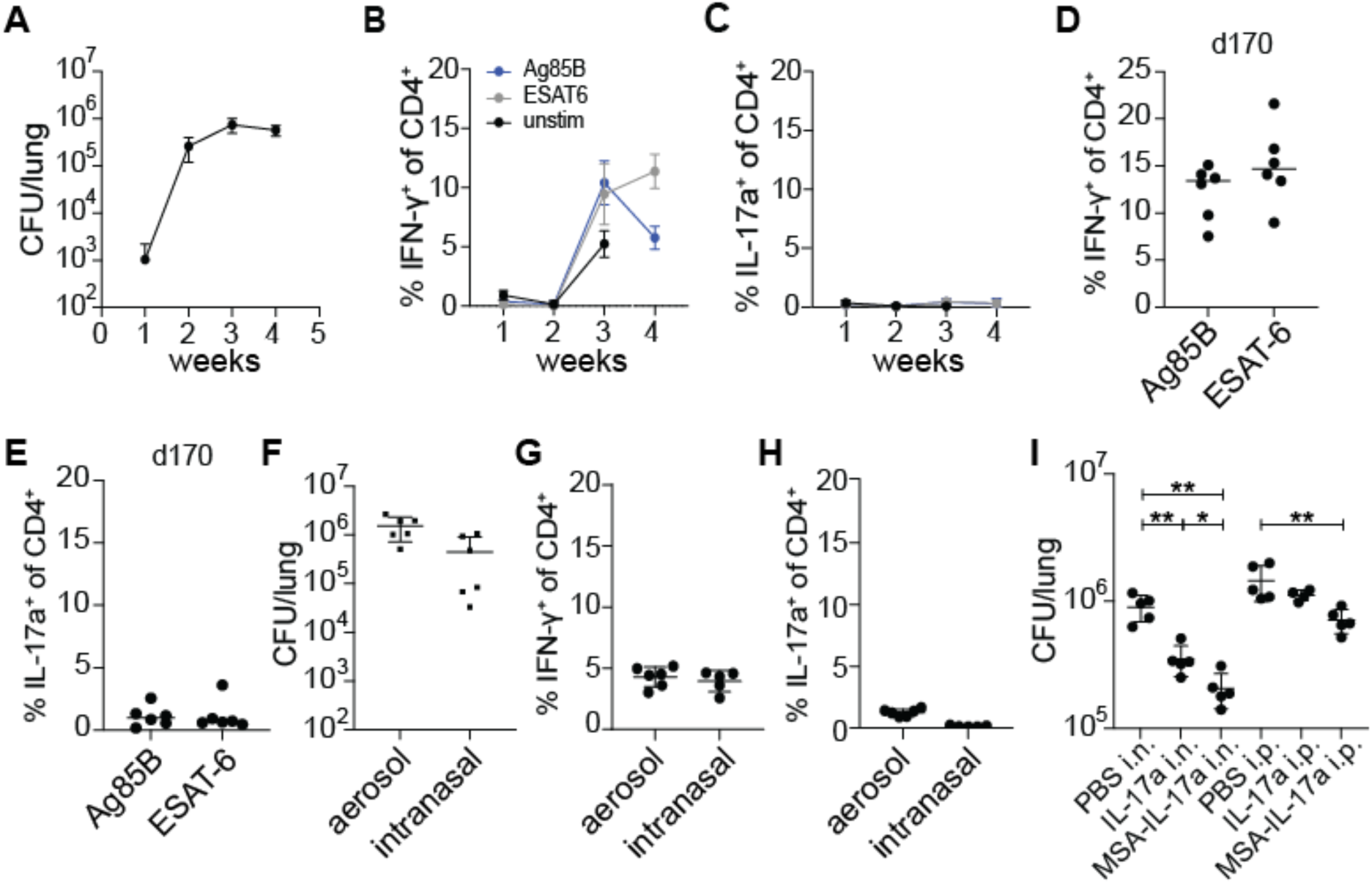
Lack of Th17 induction during WT *M. tuberculosis* infection is not due to duration or route of infection. **(A-C)** WT B6 mice were aerosol infected with WT *M. tuberculosis* Erdman strain and lungs were evaluated at 7, 14, 21, and 28 days post infection (dpi) for **(A)** bacterial burdens by CFU **(B)** IFN-γ^+^ CD4^+^ T cells and **(C)** IL-17A^+^ CD4^+^ T cells using flow cytometry. Cytokines were measured using ICS after restimulation with Ag85b or ESAT-6 peptide pools. **(D-E)** Mice were infected and analyzed at 170 dpi for **(D)** IFN-γ^+^ CD4^+^ T cells and **(E)** IL-17A^+^ CD4^+^ T cells in lungs. **(F-H)** WT B6 mice were either aerosol infected or intranasally infected with WT *M. tuberculosis* Erdman strain and evaluated 28 dpi for **(F)** bacterial burdens in lungs by CFU, **(G)** IFN-γ^+^ CD4^+^ T cells and **(H)** IL-17A^+^ CD4^+^ T cells in lungs by flow cytometry. Cytokines were measured using ICS after restimulation with ESAT-6 peptide pool. **(I)** WT B6 mice were aerosol infected with WT *M. tuberculosis* and treated with PBS control, recombinant murine IL-17A (IL-17A) or recombinant murine IL-17A fused to murine serum albumin (MSA-IL-17A) daily either using the intranasal (i.n.) or intraperitoneal (i.p.) routes between days 11 to 20 post infection. CFU in lungs was measured at 21 dpi. Results are representative of at least 2 biological replicates. For all graphs, each dot represents an individual mouse. *p<0.05, **p<0.01 (unpaired nonparametric Mann-Whitney U test).

The route of bacterial infection can impact which T helper cell types are induced. Infection via the intranasal route has been shown to elicit Th17 cells in response to infection with *Listeria monocytogenes* and Group A *Streptococcus* (12, 34). We therefore tested whether the intranasal route of infection results in a Th17 response in the lungs of Mtb infected mice. Mice were infected intranasally with ∼100 CFU of WT Mtb into the lungs as measured on day 1 after infection. After four weeks of infection, mice infected by either route had comparable levels of bacteria in the lungs (Figure 1F). At this timepoint we again observed only IFN-γ producing CD4 T cells and very few IL-17A producing CD4 T cells in the lungs (Figures 1G-H). Thus, the route of infection with Mtb does not impact the dominant Th1-biased response observed during primary infection of naïve mice with Mtb. We next investigated whether exogenous IL-17A treatment provides protection against WT Mtb infection during primary infection. WT mice aerosol infected with WT Mtb were intranasally administered with either PBS control or 1 μg recombinant murine IL-17A daily between days 11 to 20 post infection for a total of 10 doses (Figure 1I). WT mice intranasally (i.n.) administered 1 μg recombinant murine IL-17A daily were slightly more protected with ∼3-fold reduction in lung CFU compared to WT mice given vehicle control. However, we observed no effect of IL-17A when it was administered via the intraperitoneal (i.p.) route. Cytokines are known to have short half-lives in vivo. Fusing a cytokine to serum albumin can increase its half-life, permitting greater efficacy (35). We therefore fused IL-17A to murine serum albumin (MSA-IL-17A) and administered this construct either i.p. or i.n. at equimolar quantities compared to free IL-17A administered dosages. We found that i.n. delivery of this construct was more effective than delivery of IL-17A, with ∼5-fold decrease in CFU observed in the lungs. In addition, fusion to albumin allowed us to observe efficacy with i.p. administration, resulting in a ∼2-fold decrease in lung CFU. These results indicate that IL-17A can induce protective immunity to Mtb in the context of a primary infection, and that this may have potential as a therapeutic approach.

### *Tbet*^−/−^ mice are protected by a robust Th17 response during early *M. tuberculosis* infection

Because Th1 responses are known to inhibit Th17 responses (11, 13), we next tested whether Th1 differentiation masks a protective Th17 response during Mtb infection. *Tbx21*, also referred as Tbet, encodes the essential transcription factor that drives the Th1 lineage (11). Interestingly, *Tbet*^−/−^ mice lack Th1 T cells but are not as susceptible to infection with Mtb as *Ifng^−/−^*mice (5, 36). As expected, we found that *Ifng*^−/−^ mice were extremely susceptible to infection with Mtb, however *Tbet*^−/−^ had CFU in the lungs equivalent to wild type mice at 21 days after infection (Figure 2A). Only WT mice had IFN-γ producing CD4 T cells in the lungs at this timepoint (Figures 2B, 2C). Importantly, *Tbet*^−/−^ mice exhibited a robust population of IL-17A producing CD4 T cells at 3 weeks post infection, confirming that Tbet prevents Th17 differentiation during Mtb infection (Figures 2B, 2D). Importantly, while IFN-γ deficient mice exhibited a modest increase in IL-17A producing CD4 T cells (∼5-fold) compared to WT B6 mice, Tbet deficient mice had a greater 25-fold induction of IL-17A producing CD4 T cells. This demonstrates that that IFN-γ production is not the sole mechanism by which Th1 T cells limit a Th17 response during Mtb infection (Figure 2D).

**Figure 2.**
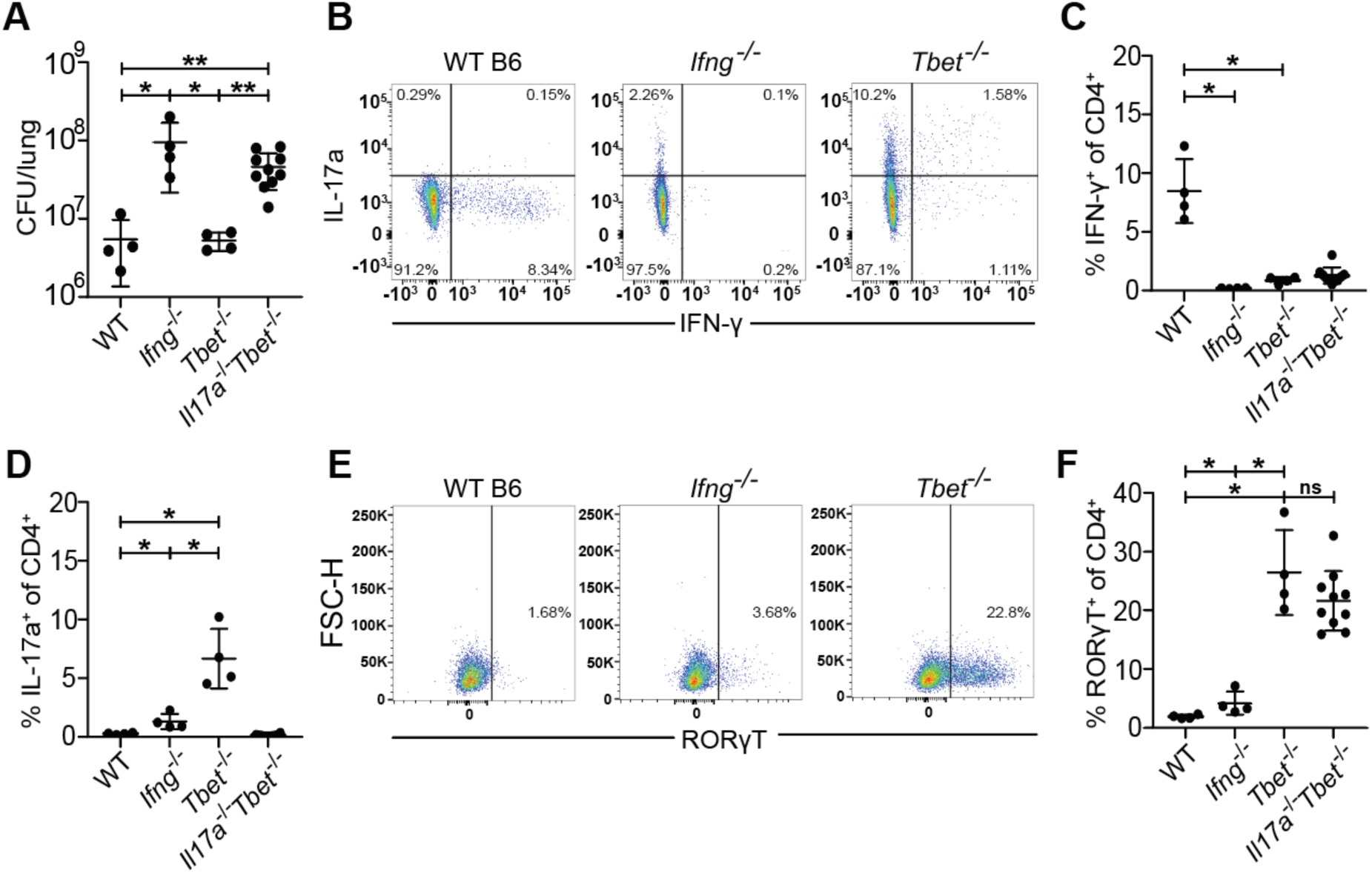
*Tbet*^−/−^ mice are protected from susceptibility to Mtb infection by the emergence of Th17 cells. WT B6, *Ifng^−/−^*, *Tbet^−/−^*, and *Il17a^−/−^Tbet^−/−^* mice were aerosol infected with WT *M. tuberculosis* Erdman strain. *Tbet^−/−^* and *Il17a^−/−^Tbet^−/−^* mice were littermate-controlled. Mice were infected with Mtb and evaluated 21 days post infection for **(A)** CFU in the lungs of indicated genotypes or **(B-F)** cellular immune responses using flow cytometry. CD4^+^ T cells were gated on single, live, MHC-II^−^, Ly6G^−^, CD3^+^, γδTCR^−^, CD4^+^, CD8a^−^ cells for analysis of IFN-γ **(B,C)**, IL-17 **(B,D)** and RORγT **(E,F)**. Results in A-F are representative of 3 independent experiments. *, p < 0.05; **, p < 0.01 (unpaired nonparametric Mann-Whitney U test).

A previous study demonstrated that Tbet constrains effective T cell responses by promoting the differentiation of terminally differentiated non-protective intravascular Th1 T cells (37). This study also used antibody blockade of IL-17A in Tbet deficient mice to conclude that IL-17A provides minimal protection in this setting (37). However, given that antibody blockade is limited by its short duration and possibly incomplete inhibition, we tested whether genetically ablating IL-17A production in Tbet mice enhanced their susceptibility to Mtb infection. We therefore infected *Tbet*^−/−^ and *Tbet*^−/−^*Il17a*^−/−^ (*Il17a^icre/icre^* referred to as *Il17a*^−/−^) littermate-controlled mice with Mtb and measured CFU and T cell responses in the lungs after 3 weeks of infection. *Tbet*^−/−^*Il17a*^−/−^mice were highly susceptible to Mtb infection with a 10-fold increase in lung CFU compared to WT and *Tbet*^−/−^ mice, comparable to the susceptibility observed in *Ifng^−/−^* mice (Figure 2A). *Tbet*^−/−^*Il17a*^−/−^ mice had significantly fewer IFN-γ producing (Figure 2C) and IL-17A producing CD4 T cells (Figure 2D). Notably, *Tbet^−/−^Il17a^−/−^* mice still generated Th17 cells, as evidenced by the presence of the Th17 master transcription factor RORγT (11), despite their inability to produce IL-17A (Figures 2E, 2F). Together, these results indicate that IL-17A production by Th17 cells confers significant protection against Mtb infection in *Tbet^−/−^* mice, at least during early stages of infection.

### *M. tuberculosis* utilizes ESX-1 and PDIM to limit Th17 response in mice

We next examined whether Mtb utilizes virulence factors to alter the host T helper response. The ESX-1 secretion system and the complex lipid PDIM are both known to influence production of cytokines known to impact Th17 differentiation (23, 24, 28, 38). Furthermore, these systems have been proposed to operate in the same virulence promoting pathway (28, 39). We therefore tested whether ESX-1 and PDIM might promote virulence in part by suppressing Th17 differentiation (Figure 3A). To test for a role of the ESX-1 secretion system, we infected mice with Δ*eccC1,* which lacks an ATPase required for ESX-1 function (ΔESX-1) (40) (Supplementary Figure S2A). We also utilized a mutant with an inactivating transposon insertion in *fadD28*, a gene required for PDIM biosynthesis (*fadD28::tn*, PDIM-lacking) (27) (Supplementary Figure S2B). As expected, ΔESX-1 and PDIM-lacking mutants were highly attenuated during infection at 3 weeks, exhibiting a 100-fold reduction in bacterial burden while complemented strains had similar CFU as WT Mtb (Supplementary Figure S3). Interestingly, ESAT-6 peptide pool stimulation of lung cells in ΔESX-1 and PDIM-lacking Mtb infected mice had an ∼5-fold reduction in %IFN-γ^+^ of CD4^+^ T cells (Th1s) (Figure 3B) and ∼2-fold increase in %IL-17A^+^ of CD4^+^ T cells (Th17s) (Figure 3C) compared to complemented strains of either mutant or WT Mtb. Similar results were seen with Ag85b peptide pool stimulation (Supplementary Figure S4A, S4B). To validate flow cytometry results we measured cytokine levels in lungs using BioLegend’s LEGENDplex kit. We found that ΔESX-1 and PDIM-lacking mutants induced less IFN-γ (Figure 3D) while also inducing ∼10x higher levels of IL-17A (Figure 3E) in the lungs compared with WT and complemented Mtb strains (Figures 3D-E). We tested if these elevated levels of Th17 cells and IL-17A contribute to the attenuation of ESX-1 and PDIM mutants by infecting littermate-controlled WT B6 and *Il17a*^−/−^ mice with either WT, ΔESX-1, and complemented ΔESX-1 Mtb, or WT, PDIM-lacking, and complemented PDIM-lacking Mtb. Under the conditions where Th17s are highly induced, mice infected with either ΔESX-1 or PDIM-lacking Mtb, the *Il17a*^−/−^ mice had ∼3-5 fold higher CFU than WT mice (Figures 3F-G). These results indicate that the induction of Th17s is not dependent on the attenuation of Mtb in general, but instead Mtb utilizes ESX-1 and PDIM to suppress the induction of a Th17 response that enhances protection against Mtb infection.

**Figure 3.**
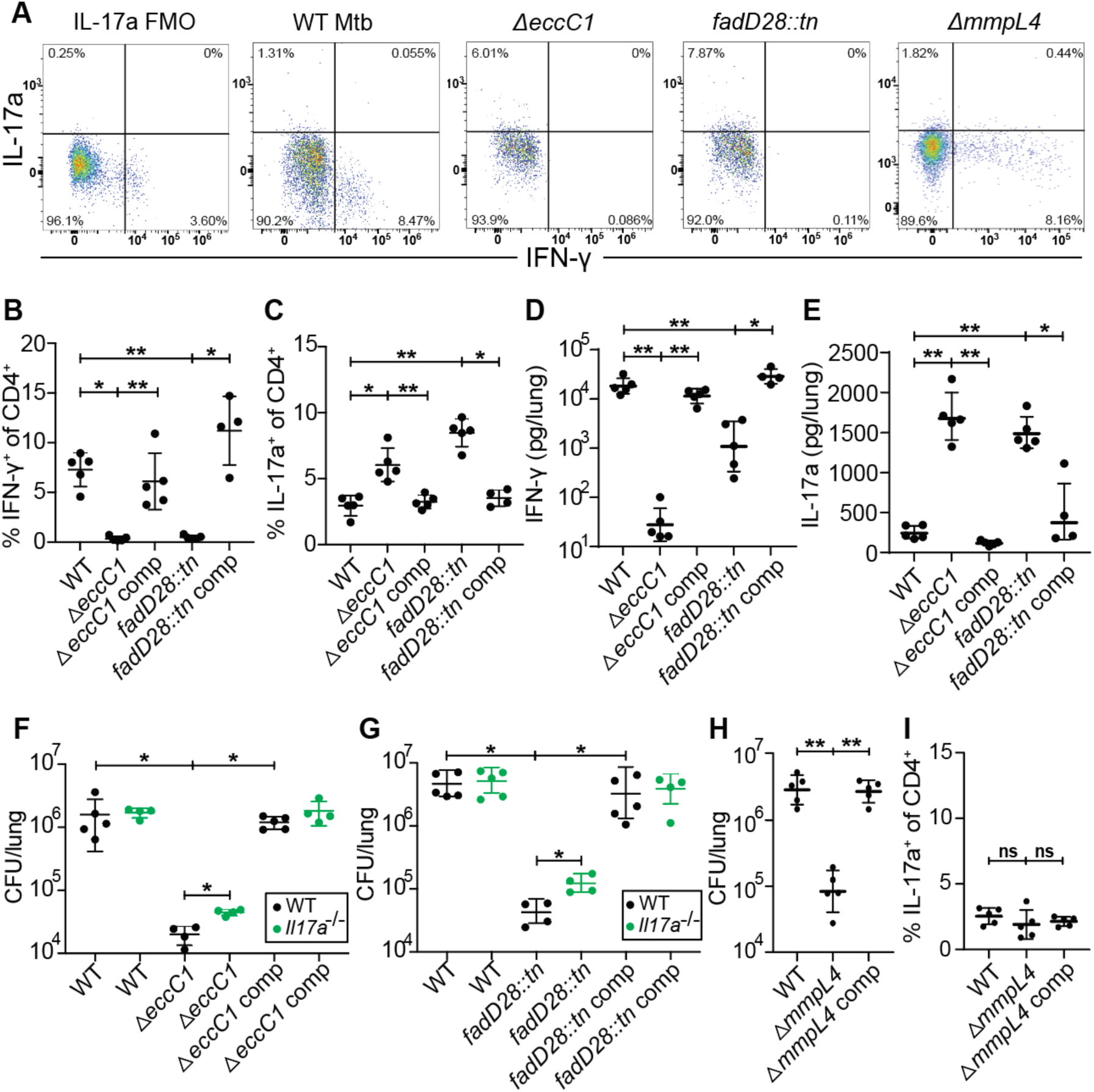
*M. tuberculosis* ESX-1 and PDIM virulence factors suppress Th17 response in mice. WT B6 mice were aerosol infected with indicated strains and evaluated at 21 days post infection. **(A)** Representative flow cytometry plots (cells gated on single, live, MHC-II^−^, Ly6G^−^, CD3^+^, γδTCR^−^, CD4^+^, CD8a^−^) of IFN-γ^+^ and IL-17A^+^ CD4^+^ T cell populations. Mice were sacrificed and evaluated with ESAT-6 peptide pool stimulated lung homogenates for **(B)** IFN-γ^+^ CD4^+^ T cells and **(C)** IL-17A^+^ CD4^+^ T cells by flow cytometry. Mice were sacrificed and lung homogenates were evaluated with BioLegend LEGENDplex for **(D)** IFN-γ and **(E)** IL-17A protein concentrations. **(F-G)** Littermate-controlled WT B6 and *Il17a^−/−^* mice were aerosol infected with indicated strains and sacrificed at 21 days post infection with lung homogenate plated for CFU. **(H-I)** Mice were infected with WT, *mmpl4* mutant, or complemented *mmpl4* mutant Mtb and evaluated for **(H)** day 21 CFU and **(I)** IL-17A^+^ CD4^+^ T cells by flow cytometry. Results are representative of 2 independent experiments. *, p < 0.05; **, p < 0.01 (Mann-Whitney U test).

To ensure that the increase in Th17s and IL-17A was not simply a consequence of infection with any attenuated Mtb mutant, we evaluated responses during infection with MmpL4 deficient (Δ*mmpL4*) bacteria. MmpL4 is a transmembrane transporter with proposed siderophore secretion activity (41) with no known association with the function of ESX-1 or secretion of PDIM (27). Δ*mmpL4* Mtb-infected mice had ∼1.5 logs lower lung CFU than WT or complemented Mtb-infected mice (Figure 3H), a similar degree of attenuation as we observed with ESX-1 and PDIM mutants (Supplemental Figure S3). However, the Δ*mmpL4* Mtb-infected mice induced Th1 response that was indistinguishable from WT and complemented Mtb-infected mice (Supplemental Figure S5) while not inducing greater numbers of Th17s compared to WT or complemented Mtb-infected mice (Figure 3I). Thus, the increase in Th17s we observed results from an ESX-1/PDIM specific mechanism.

### Type I interferons boost Th1 induction and do not impact Th17 induction

As ESX-1 and PDIM are responsible for the host type I interferon response to Mtb infection (24, 28), and type I IFN is known to suppress Th17 responses (42), we investigated the impact of type I interferon signaling on Th17 induction during Mtb infection. We examined the effect of type I IFN both in WT B6 mice and in *Sp140^−/−^* mice that are highly susceptible to infection because of an excessive type I IFN response to infection (43). To abrogate type I IFN in each background we used mice deficient for the type I IFN receptor IFNAR. WT, *Ifnar1*^−/−^, *Sp140*^−/−^, *Sp140*^−/−^*Ifnar1*^−/−^, *Ifng*^−/−^, and *Tbet*^−/−^ mice were aerosol infected with WT Mtb. As expected, both *Sp140*^−/−^ and *Ifng*^−/−^ mice had up to 10-fold higher CFU than WT mice while *Ifnar1*^−/−^, *Sp140*^−/−^*Ifnar1*^−/−^, and *Tbet*^−/−^ mice were as resistant as WT mice at 3 weeks post infection (Figure 4A). *Sp140*^−/−^ mice had an almost 2-fold increase in %IFN-γ^+^ of CD4^+^ (Th1) T cells compared to WT mice, *Ifnar1*^−/−^had a 2-fold reduction in %IFN-γ^+^ of CD4^+^ (Th1) T cells compared to WT mice, *Sp140*^−/−^*Ifnar1*^−/−^mice had a slightly stronger 4-fold reduction in %IFN-γ^+^ of CD4^+^ (Th1) T cells compared to *Sp140*^−/−^ mice (Figures 4B-D). Importantly, deletion of IFNAR had no impact on Th17 induction in either WT or *Sp140*^−/−^ mice, although we observed the expected increase in Th17 cells in Tbet deficient animals (Figure 4E). These results indicate that type I interferon signaling only impacts the induction of Th1s and does not influence Th17 induction, suggesting that induction of type I IFN by ESX-1/PDIM is not the mechanism by which these virulence factors suppress Th17 differentiation.

**Figure 4.**
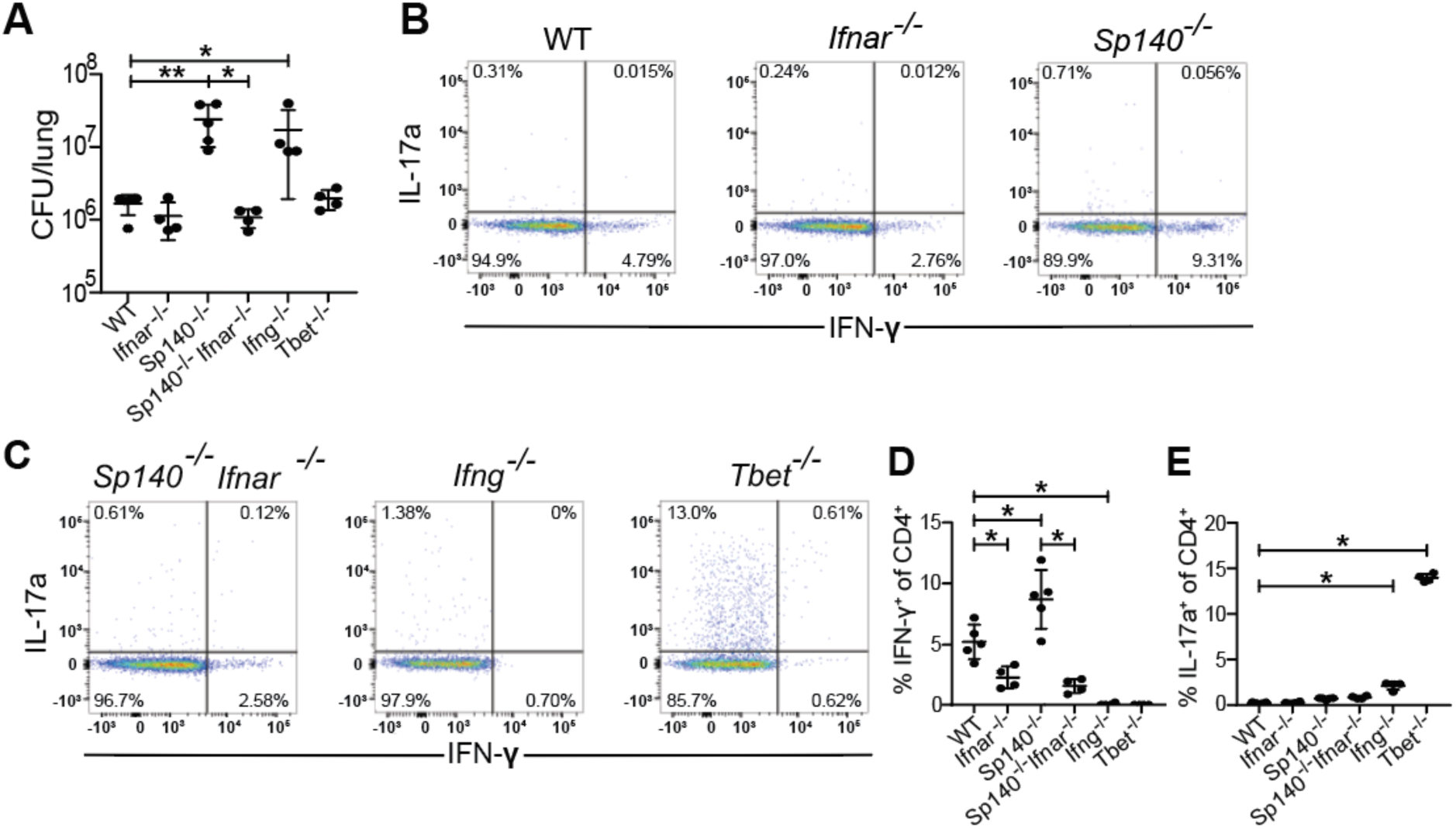
Type I interferon signaling boosts Th1 induction and does not impact Th17 induction. WT B6, *Ifnar^−/−^*, *Sp140^−/−^*, *Sp140^−/−^Ifnar^−/−^*, *Ifng^−/−^*, and *Tbet^−/−^* mice were aerosol infected with WT *M. tuberculosis* Erdman strain. **(B, C)** Representative flow cytometry plots (cells gated on single, live, MHC-II^−^, Ly6G^−^, CD3^+^, γδTCR^−^, CD4^+^, CD8a^−^) of IFN-γ^+^ and IL-17A^+^ CD4^+^ T cells of these populations. 21 days post infection, lung homogenates were measured for **(A)** CFU and analyzed via flow cytometry for **(D)** IFN-γ^+^ CD4^+^ T cells and **(E)** IL-17A^+^ CD4^+^ T cells. Results in A, D, and E are representative of 2 independent experiments. *, p < 0.05; **, p < 0.01 (unpaired nonparametric Mann-Whitney U test).

### ESX-1 and PDIM alters conventional dendritic cell cytokine signaling in the mediastinal lymph node

Previous work suggested that ESX-1 mutants fail to suppress expression by macrophages of IL-12 p40, a subunit shared between IL-12 and IL-23 (24). Because IL-23 promotes Th17 responses, and ESX-1 and PDIM are responsible for the lack of a Th17 response during Mtb infection, we next investigated if ESX-1 and PDIM suppress IL-23 production in bone-marrow-derived macrophages (BMDMs) during in-vitro infection. ESX-1 and PDIM deficient strain both elicited a strong IL-23 response measured by IL-23 ELISA (Figure 5A). We then investigated if ESX-1 and PDIM suppress IL-23 production by dendritic cells during in vivo infection in the draining (mediastinal) lymph nodes. We infected mice with ∼100 CFU via the aerosol route and harvested mediastinal lymph nodes at 21 days post infection. We examined IL-12 and IL-23 production by conventional dendritic cells using ICS. To clearly distinguish between levels of these two cytokines we used ICS for the cytokine specific subunits: IL-12 p35 and IL-23 p19.

**Figure 5.**
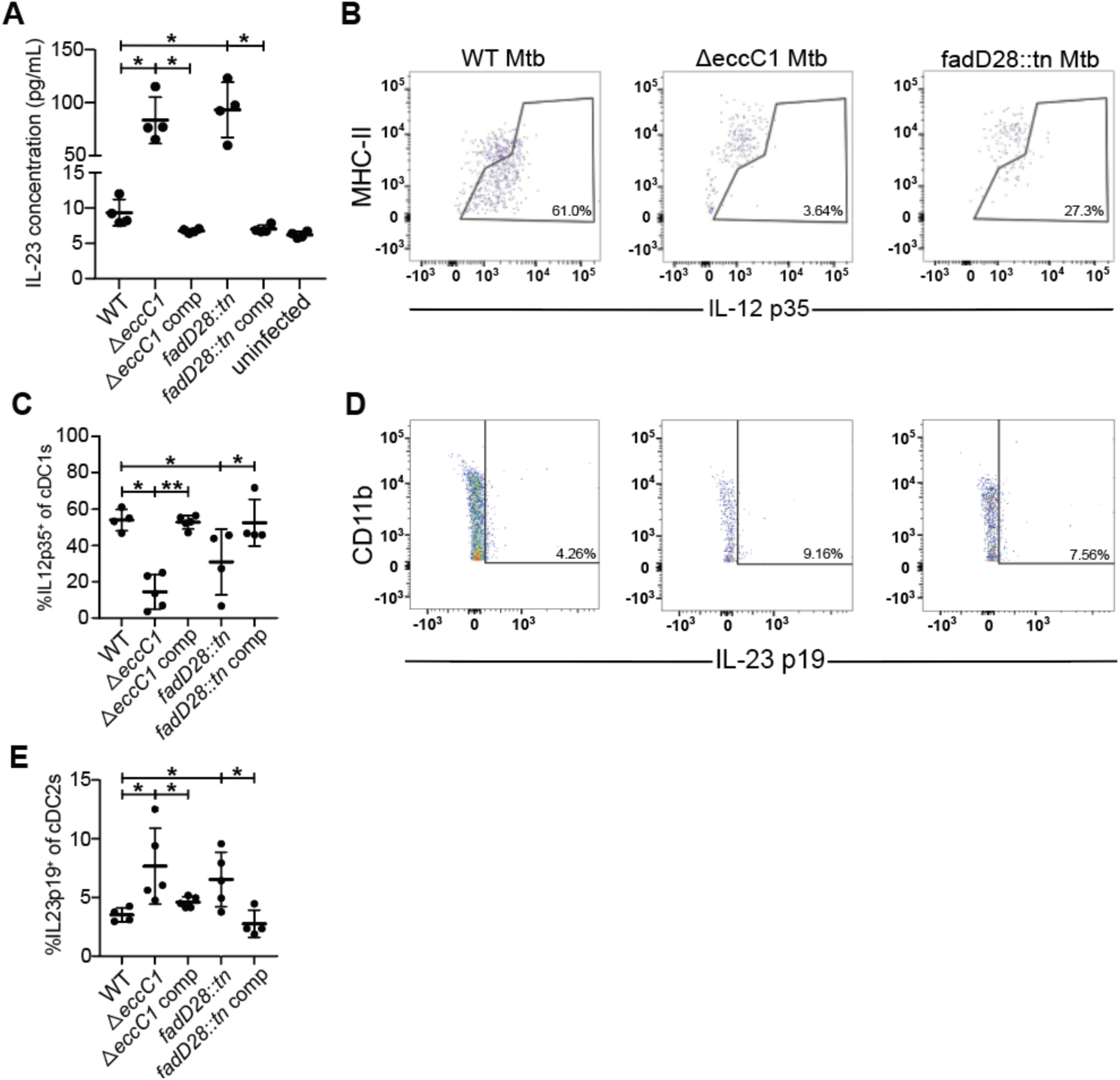
ESX-1 and PDIM alter conventional dendritic cell cytokine signaling in the mediastinal lymph node. WT B6 bone-marrow-derived macrophages were infected with designated Mtb Erdman strains at MOI = 10 or uninfected for 24 hours with **(A)** IL-23 ELISA taken from cultured supernatants. WT B6 mice were aerosol infected with indicated strains and analyzed at 21 days post infection **(C,E)** Flow cytometry schematic for gating of **(B)** cDC1 (gated on single, live, CD3^−^, CD19^−^, CD11c^+^, MHC-II^+^, XCR1^+^, CD11b^−^) and **(D)** cDC2 (gated on single, live, CD3^−^, CD19^−^, CD11c^+^, MHC-II^+^, XCR1^−^, CD11b^+^) populations of mediastinal lymph nodes with representative flow cytometry plots of IL-12 p35^+^ and IL-23 p19^+^ gating of cDC1 and cDC2 populations respectively. 21 days post infection, mediastinal lymph nodes from WT B6 mice were aerosol infected with either WT, ΔeccC1 (ΔESX-1), complemented ΔeccC1::eccC1, transposon-inserted fadD28::tn (PDIM-lacking),or complemented transposon-inserted fadD28::tn + pfadD28 *M. tuberculosis* Erdman strain were extracted and analyzed for IL-12 p35^+^ cDC1s **(C)** and IL-23 p19^+^ cDC2s **(E)** by flow cytometry. Results in A, C, and E are representative of 2 independent experiments. *, p < 0.05; **, p < 0.01 (unpaired nonparametric Mann-Whitney U test).

First, we found that type 1 conventional dendritic cells (cDC1s), known to be specialized for IL-12 production, produce robust levels of IL-12 p35 during infection with WT Mtb (Figures 5B, 5C). However, both ESX-1 and PDIM deficient strains elicited significantly lower levels of IL-12 p35 from these cells (Figures 5B, 5C), consistent with the lower bacterial burdens in the lungs of mice infected with these mutants (Supplementary Figure S3). During infection with wild-type bacteria, we observed very little IL-23 p19 production by type 2 conventional dendritic cells (cDC2s) in the draining mediastinal lymph node (Figures 5D, 5E). Importantly, ESX-1 and PDIM-lacking Mtb infection resulted in an increase in the percentage of cDC2s that produce IL-23 p19 in mediastinal LNs (Figures 5D, 5E). We saw very little to no IL-23 p19 expression in cDC1s and IL-12 p35 expression in cDC2s with no differences between WT or ESX-1 and PDIM deficient strains (Supplementary Figure S6). These results indicate that the ESX-1 and PDIM virulence factors impact T helper cell subsets observed in the lungs by promoting IL-12, a Th1-polarizing cytokine, and limiting IL-23, a Th17-polarizing cytokine, expression in cDC1s and cDC2s respectively in the draining mediastinal lymph node which may impact naïve T cell differentiation.

## Discussion

The rise of multidrug-resistant and extensively drug-resistant strains, combined with a limited pipeline of new drugs, underscores the urgent need for effective vaccines. Achieving the World Health Organization/STOP TB Partnership’s target of TB elimination by 2050 will not be possible without a highly efficacious vaccine (44). Unlike most successful vaccines, which protect through neutralizing antibodies, TB immunity primarily depends on T-cell–mediated responses, for which no clear vaccine strategy exists. Defining protective immune mechanisms is therefore critical—not only for guiding vaccine design but also for advancing immune-based therapies. Equally important is understanding how *M. tuberculosis* evades or suppresses these responses, as such insights can inform both vaccine development and therapeutic interventions.

We sought to understand why Mtb infection does not elicit Th17 production robustly during primary infection. We found that this is independent of duration and route of infection but instead results from an overly robust Th1 response, which suppresses Th17 development, and upon the virulence factors ESX-1 and PDIM. Perhaps Mtb uses ESX-1 and PDIM to allow other virulence factors like Hip1 to access the host cytosol, alter host cellular signaling, and impair dendritic cell expression of IL23 p19, CD40, Jagged, and DLL4 shown to be important for dendritic cell-mediated differentiation of naïve T helper cells into Th17s (45–48). It has been long established that Th1 cells are important for control of Mtb infection, yet there have also been hints in the literature that Th1 cells are not simply protective. Firstly, excessive IFN-γ production is harmful, driving tissue destruction in the lungs (49), which may benefit bacteria by promoting necrosis and transmission. Interestingly, the majority of T cell antigens are highly conserved across Mtb (50), suggesting that dominant Th1 cell responses may confer some benefit to Mtb. Indeed, a recent study suggested that while common antigens are likely to drive Th1 differentiation, rare variable antigens are more likely to drive Th17 differentiation in Mtb infected patients (51). Thus, it is feasible that Mtb promotes excessive Th1 differentiation in part to prevent potentially protective Th17 cells from emerging.

Given that ESX-1 and PDIM mutants elicit type I IFN, and type I IFN suppresses Th17 cells, it is perhaps surprising that type I IFN does not explain the suppression of Th17 we observe during early infection. Type I IFN is clearly detrimental to Mtb infection, as deletion of IFNAR rescues the susceptibility of mice that exhibit type I IFN driven disease (24, 43). However, type I IFN is known to have many immunosuppressive functions that could explain its detrimental role in Mtb infection (52). Many cytokines are known to participate in the differentiation of Th17 cells including TGF-β, IL-6 and IL-23. Importantly, IL-23 is essential for maintaining the Th17 lineage, while TGF-β and IL-6 are critical for its induction. The crucial role of IL-23 differentiation and maintenance of Th17 cells in vivo is well established (53, 54). IL-23 supports IL-17 production by upregulating IL-17 itself, RORγt and other pro-inflammatory cytokines (55). Our in vivo data demonstrating increased IL-23 production by dendritic cells in ESX-1 mutant infections is consistent with the old observation that ESX-1 suppresses IL-12 p40 in vitro. Future work will focus on elucidating the mechanism of Mtb suppression of IL-23. It is also possible that ESX-1/PDIM suppress Th17 differentiation by promoting Th1 differentiation, however we do not have data that supports this model at this time. Finally, the work in this study adds to the growing evidence that ESX-1 and PDIM are phenotypically linked. Unlike what has been demonstrated in the H37Rv strain (28), we do not find that PDIM mutants have a defect in ESX-1 function or that ESX-1 mutants have any changes in PDIM production or secretion. This could be a strain specific difference, especially as it is known that H37Rv has a unique point mutation that causes downregulated ESX-1 function relative to Erdman and most clinical strains (56). In the future, it will be important to establish how ESX-1 and PDIM function are linked functionally. Secretion of the ESX-1 substrate ESAT-6 is generally used as a proxy for ESX-1 function. It is possible that PDIM loss has more subtle effects on ESX-1 function that are not reflected by ESAT-6 secretion levels.

Because primary infection does not reliably elicit Th17 T cells, it has been difficult to study the function of these cells. A method for reliably inducing Th17s in mice during infection would facilitate studying the role of Th17-mediated protection. We show here that mice lacking Tbet respond to WT Mtb infection with a Th17 response and provide protection mediated by IL-17A during early infection. Thus, *Tbet*^−/−^ mice can be used to interrogate mechanisms underlying Th17 mediated protection during primary Mtb infection. High Th17 plasticity permits Th17s to express a wide variety of T helper subset profiles that can range from pro-inflammatory Th17.1s that co-express IFN-γ and IL-17A to anti-inflammatory Tr1-like Th17s that co-express IL-10 and IL-17A (57). Moving forward, it will be important to delineate in depth which Th17 subsets and which mechanisms downstream of IL-17 receptor signaling are important for controlling Mtb infection. Importantly, we show that exogenous administration of recombinant IL-17A leads to a decrease in CFU in Mtb infected lungs by up to ∼5x. Our data suggests that a limiting factor for efficacy of recombinant IL-17A is its short half-life. Future efforts focused on methods for improving the stability of this cytokine in vivo could lead to more dramatic reductions in bacterial burdens.

The live vaccine strain Bacille Calmette-Guerin (BCG) was developed by extensive laboratory passage of *Mycobacterium bovis*, resulting in attenuation of virulence (58, 59). This attenuation makes BCG a very safe vaccine. However, although BCG is effective in preventing severe tuberculosis in children, it is not sufficiently effective against adult pulmonary tuberculosis to make a significant impact on TB cases worldwide. A cause of the reduced virulence of BCG is the loss of multiple genes associated with the ESX-1 type VII secretion system. To create a live vaccine that is safe yet more effective than BCG, scientists engineered MTBVAC, a vaccine candidate currently in phase 2b trials. Unlike BCG, MTBVAC is derived from Mtb. MTBVAC is attenuated by mutation of the ESX-1 secretion system and a mutation that eliminates PDIM (60). Our insights into how ESX-1 and PDIM suppress effective immune responses could promote understanding of how MTBVAC provides protective immunity. Indeed, MTBVAC has been demonstrated to elicit mucosal-homing-marker expressing Th17s ideal for better control of Mtb infection (61). In addition, understanding the role and function of Th17 cells more generally can help promote development of the next generation of effective Mtb vaccine candidates.

## Materials and Methods

### Ethics statement

All procedures involved with the use of mice were approved by the University of California, Berkeley’s Institutional Animal Care and Use Committee (protocol 2015-09-7979). All protocols conform to federal regulations, the National Research Council Guide for the Care and Use of Laboratory Animals, and the Public Health Service Policy on Humane Care and Use of Laboratory Animals.

### Bacterial Culture

*Mycobacterium tuberculosis* strain Erdman was grown in Middlebrook 7H9 liquid medium supplemented with 10% oleic acid, albumin, dextrose, and catalase (OADC) (BD 212351), 0.4% glycerol, and 0.05% Tween 80 or on solid 7H10 agar plates supplemented with 10% OADC and 0.4% glycerol. Frozen stocks of *M. tuberculosis* were made from a single culture and used for all experiments.

### Mice

C57BL/6J (no. 000664), *Ifng*^−/−^ (B6.129S7-*Ifng*^tm1Ts^/J, no. 002287), *Tbet*^−/−^ (*Tbx21*^−/−^, B6.129S6-*Tbx21*^tm1Glm^/J, no. 004648), and *Il17a*^icre/icre^ (*Il17a*^−/−^, B6.129(SJL)-*Il17a*^tm1.1(icre)Stck^/J, no. 035717) mice were obtained from The Jackson Laboratory and bred in-house. *Ifnar1*^−/−^, *Sp140*^−/−^, and *Sp140*^−/−^*Ifnar1*^−/−^ mice were obtained from the Vance laboratory (UC Berkeley) and bred in-house. Mice were age matched for all experiments.

### Aerosol challenge experiments with *M. tuberculosis*

Mice between the age of 8 to 12 weeks old were infected via the aerosol route with *M. tuberculosis* strain Erdman using a nebulizer and full-body inhalation exposure system (Glas-Col, Terre Haute, IN). A total of 9 mL of diluted aerosol stock in MilliQ water was loaded into the nebulizer to deliver 30-100 bacteria per mouse as measured by whole lung CFU 1 day post infection.

### Intranasal challenge experiments with *M. tuberculosis*

Mice between the age of 8 to 12 weeks old were infected with 40 uL of *M. tuberculosis* strain Erdman diluted in PBS administration for mice to inhale 30-100 bacteria per mouse as measured by whole lung CFU 1 day post infection.

### Flow Cytometry

Whole lungs were homogenized using GentleMACS C tubes (Miltenyi Biotec 130-093-237) in complete RPMI-1640 media supplemented with 10% FBS, 1 mM sodium pyruvate (Sigma Aldrich S8636), 0.01 mM HEPES, 1% Glutamax (Thermo Scientific 35050061), 1% non-essential amino acids (Gibco 11140-050), and 55 μM 2-mercaptoethanol (Gibco 21985-023) containing Liberase TM (Sigma Aldrich 05401119001) and DNase I (Sigma Aldrich 11284932001). Cells were strained through 70 micron strainer (Corning 352350). Cells were treated with protein transport inhibitor cocktail mix (Invitrogen 00-4980-03) and stimulated with vehicle control, Mtb Ag85b peptide pool (Peptides&elephants LB02088), Mtb esat6 peptide pool (Peptides&elephants LB02113), or PMA and ionomycin (Invitrogen 00-4970-93) for 5 hours at 37 °C, 5% CO_2_. Cells were stained with live/dead stain (ThermoFisher L23101), Fc block (Biolegend 101320), and surface stained with antibodies for CD3, Ly6G, CD4 (BD 740268, 551460, and 569182 respectively), MHC-II, pan-γδTCR, and CD8a (Biolegend 107605, 118124, and 100714 respectively) in Brilliant stain buffer (BD 566349). Cells were fixed and permeabilized with Foxp3 staining kit (Invitrogen 00-5523-00) before intracellular staining for RORγT, Foxp3 (BD 564723 and 562996 respectively), Tbet, IFN-γ, and IL-17A (Biolegend 644817, 505814, and 506903 respectively). Data was collected using a BD Symphony A3 flow cytometer (BD) and analyzed using FlowJo Software (BD).

Mediastinal lymph nodes were homogenized using complete RPMI-1640 containing Liberase TM (Sigma Aldrich 05401119001) and DNase I (Sigma Aldrich 11284932001) with sterile plunger of 3 mL syringe to pass dissociated tissue through 100 micron cell strainer (Corning 352360). Cells were stained with live/dead stain (ThermoFisher L34966), Fc block (Biolegend 101320) and surface stained with antibodies for XCR1, CD11b, CD11c, Ly6G, MHC-II, CD3, CD19 (Biolegend 148220, 101261, 117318, 127627, 107606, 100233, and 115545 respectively) and Ly6C and CD64 (BD 755198 and 569507 respectively) in Brillant stain buffer (BD 566349). Cells were fixed and permeabilized with Foxp3 staining kit (Invitrogen 00-5523-00) before intracellular staining for IL-12 p35 (Invitrogen MA5-23559) and IL-23 p19 (BD 565317). Data were collected using a BD Symphony A3 flow cytometer (BD) and analyzed using FlowJo Software (BD).

For Legendplex, whole lungs were homogenized using 70 micron cell strainer (Corning 352350) and plunger of 3 mL syringe into complete RPMI. Single cell suspensions were pelleted at 1200 rpm for 5 minutes at 4 °C. Supernatants were collected for LEGENDplex with pre-made mouse inflammation panel (13-plex) (Biolegend 740446). Prior to final wash and running on flow cytometer, beads were fixed with 4% PFA and plate transferred to BSL1. Data was collected using a BD Symphony A3 flow cytometer (BD) and analyzed using LEGENDplex Data Analysis Software Suite (Biolegend).

### Lung homogenate protein concentration measurements

Whole lungs were homogenized in complete RPMI-1640 media supplemented with 10% FBS, 1 mM sodium pyruvate (Sigma Aldrich S8636), 0.01 mM HEPES, 1% Glutamax (Thermo Scientific 35050061), 1% non-essential amino acids (Gibco 11140-050), and 55 μM 2-mercaptoethanol (Gibco 21985-023) and meshed through 70 micron cell strainer (Corning 352350) with plunger of 5 mL syringe. Cells were pelleted at 1,500 rcf for 5 minutes and supernatants were collected for measuring protein concentrations with BioLegend LEGENDplex mouse inflammation panel (13-plex) (Biolegend 740446).

### Exogeneous IL-17A administration

Mice were intranasally administered 1 μg endotoxin-free recombinant murine IL-17A (R&D systems, 7956-ML-100/CF) in 40 μL sterile PBS daily or 40 μL of vehicle control daily, or intraperitoneally administered 2 μg endotoxin-free recombinant murine IL-17A in 100 μL sterile PBS daily or 100 uL of vehicle control dailiy from days 11 to 20 post aerosol *M. tuberculosis* challenge.

MSA-IL-17A was generated by fusing a (GGGGS)3 linker between the C-terminus of mature murine serum albumin and N-terminus of mature murine IL-17A. MSA-IL-17A was purchased from Genscript USA, Inc. in endotoxin-free pH 7.2 PBS. Mice were intranasally administered 5.54 μg MSA-IL-17A (equimolar to 1 μg recombinant murine IL-17A) in 40 μL sterile PBS daily or 40 μL vehicle control, or intraperitoneally administered 11.09 μg MSA-IL-17A (equimolar to 2 μg recombinant murine IL-17A) in 100 μL PBS or 100 μL vehicle control daily between days 11 to 20 post aerosol *M. tuberculosis* challenge.

### Bone-marrow-derived macrophage infection

WT C57BL/6J bone-marrow-derived macrophages (BMDMs) were incubated on tissue-culture plates for 48 hours prior to infection at 37 °C, 5% CO_2_. BMDMs were spin-fected at 1200 rpm for 10 minutes at room temperature for an MOI of 10. BMDMs were washed with 1x PBS (pH 7.4) and incubated with fresh BMDM media at 37 °C, 5% CO_2_ for 24 hours. 24 hours post infection, cultured supernatants were extracted for IL-23 ELISA (Biolegend 433704).

### Statistical Analysis

All data was analyzed using FlowJo v10 (FlowJo, LCC) and/or Prism 10 (GraphPad Software). All graphs were analyzed using Mann-Whitney U-test. Significant values are symbolized with asterisks of * < 0.05, ** < 0.01, and ns for non-significant.

## Acknowledgements

We thank Russell Vance’s lab for the kind gifts of *Sp140*^−/−^, *Ifnar1*^−/−^, and *Sp140*^−/−^*Ifnar1*^−/−^ mice. We thank Jeff Cox’s lab for the kind gifts of ΔESX-1 (Δ*eccC1*), complemented ΔESX-1, PDIM-lacking (Δ*fadD28*), complemented PDIM-lacking, ΔMmpL4 (Δ*mmpL4*), and complemented MmpL4 Mtb Erdman strains. We thank Helia Samani, Scott Espich, Vicky Wu, Malia Wilson, and Cielo Kwon for assistance in mouse colony maintenance. We thank members of the Stanley, Cox, and Vance laboratories for helpful discussions. Funding was from NIH P01 AI063302 Project 4 to SAS and NIH 1R01AI113270-01A1 to SAS.

---

**Figure S1.**
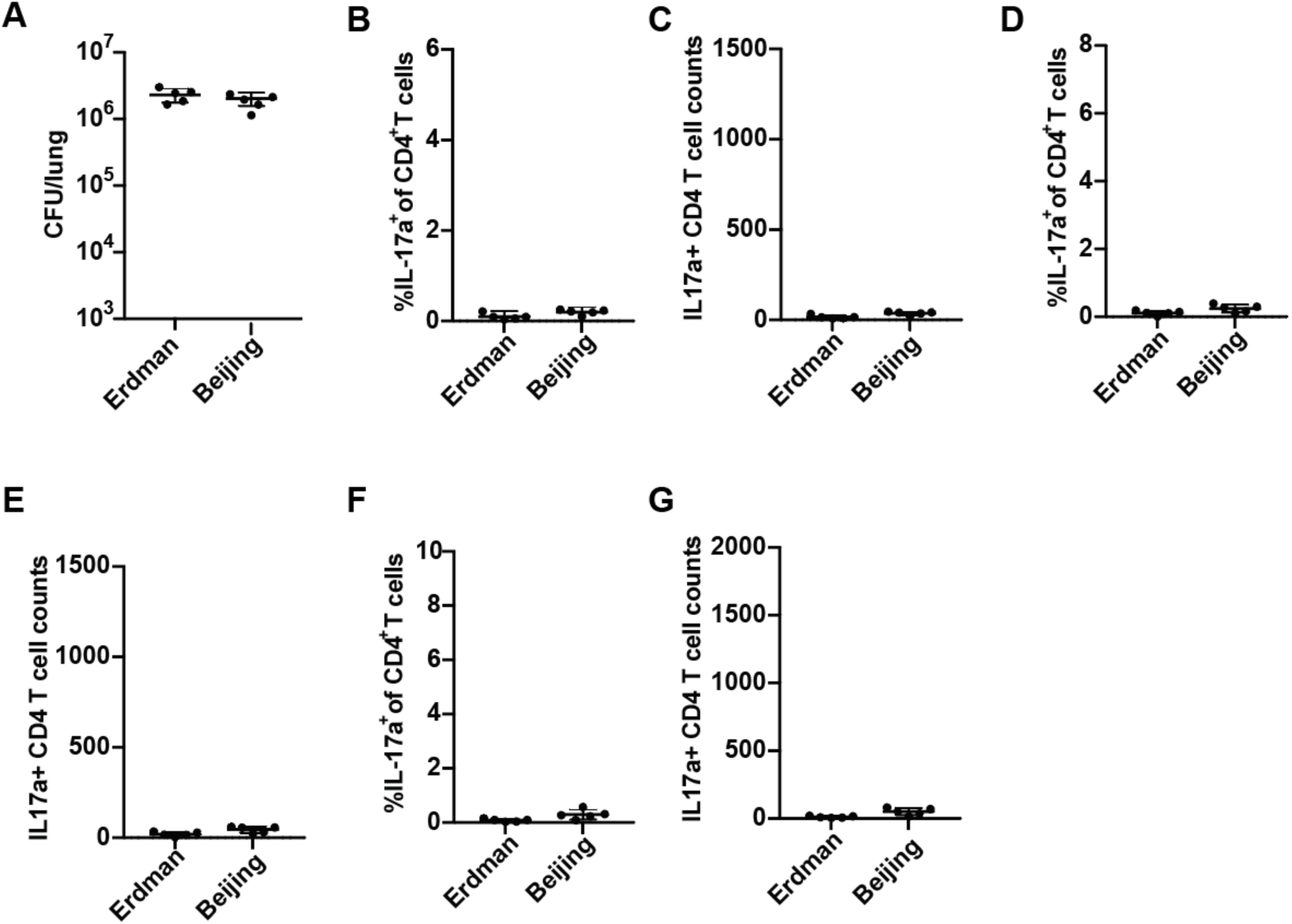
Mice were infected with either the Erdman or Beijing strain of Mtb via the aerosol route and evaluated at 21dpi for **(A)** CFU in the lungs or **(B-G)** IL-17A^+^ CD4^+^ T cells in the lungs. **(B,C)** unstimulated, **(D,E)** restimulated with Antigen 85B peptide, **(F,G)** restimulated with ESAT-6 peptide. Representative experiment of 2.

**Figure S2:**
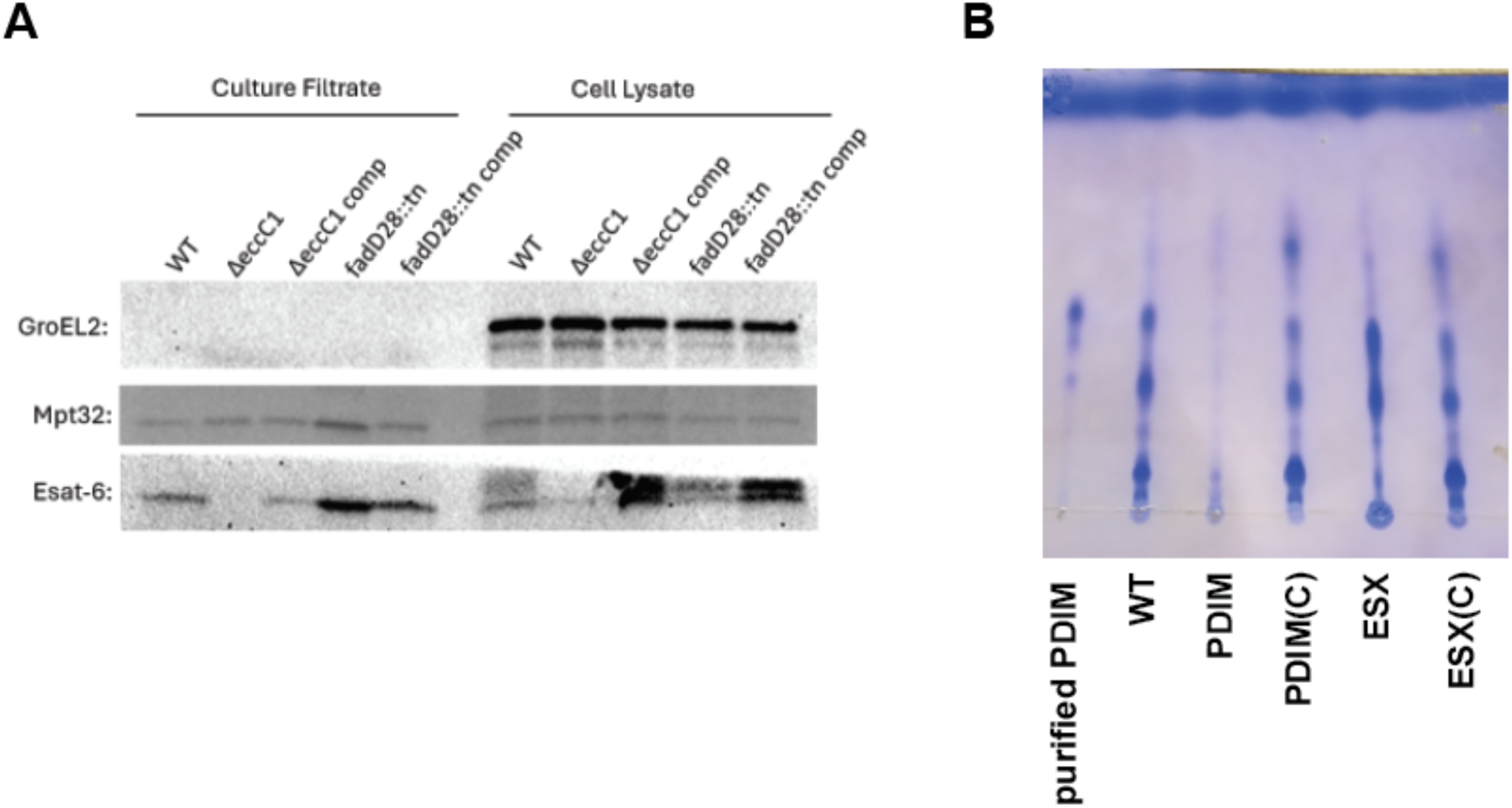
Mtb Erdman PDIM mutants secrete ESAT-6 and ESX-1 mutants produce PDIM. **(A)** WT, Δ*eccC1*, complemented Δ*eccC1::eccC1* (Δ*eccC1* comp), *fadD28::tn*, or *fadD28::tn* + p*fadD28* (fadD28::tn comp) *M. tuberculosis* Erdman strain were cultured in Sauton’s complete media without Tween-80 for 5 days. Cell lysate and culture supernatant were processed through SDS-PAGE and Western blot analysis. Anti-ESAT-6 antibody was used for detecting ESAT-6. Anti-Mtb GroEL2 antibody (BEI Resources NR-13657) was used as a loading control for the cell lysate fraction. Anti-Mtb Mpt32 antibody (BEI Resources NR-13807) was used as a loading control for the culture filtrate. Results shown are from 1 experiment representative of 2 independent experiments. **(B)** WT, Δ*eccC1* (ESX), complemented Δ*eccC1::eccC1* (ESX(C), *fadD28::tn* (PDIM), or *fadD28::tn* + p*fadD28* (PDIM(C)) *M. tuberculosis* Erdman strain were cultured in 7H9 media. Outer leaflet of mycomembrane was removed via hexanes wash, concentrated and separated on TLC plate with 98:2 petroleum ether:acetone mobile phase. TLC plate was stained with 0.2% Coomassie blue in 20% methanol. Results shown are from 1 experiment representative of 2 independent experiments.

**Figure S3.**
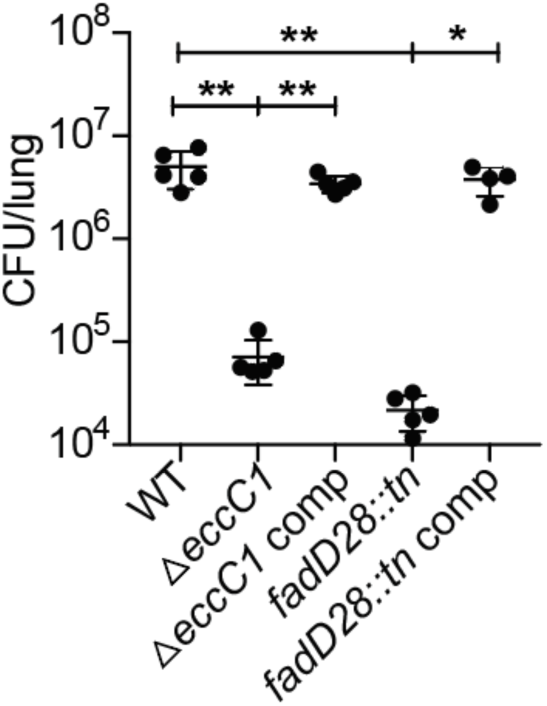
Attenuation of ESX-1 and PDIM mutants at 21 days post infection. WT B6 mice were aerosol infected with either WT, Δ*eccC1,* Δ*eccC1::eccC1*, *fadD28::tn*, or *fadD28::tn* + p*fadD28 M. tuberculosis* Erdman strain. 21 days post infection, mice were sacrificed, and CFU were enumerated from lung lysates.

**Figure S4:**
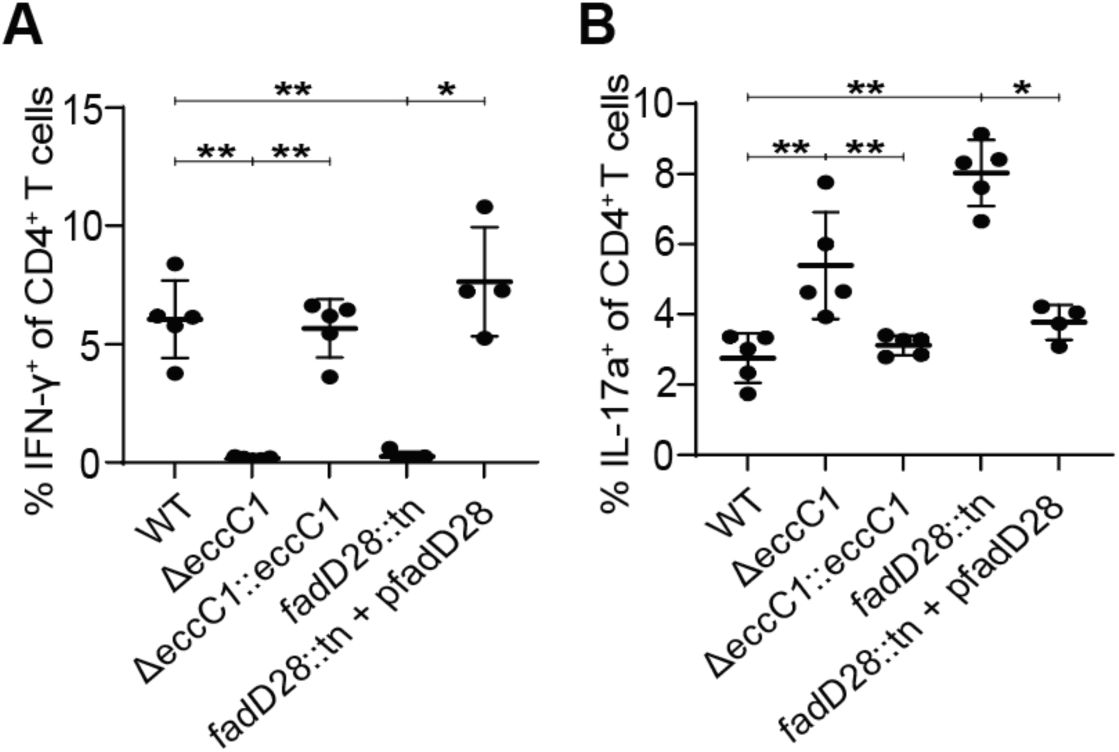
Antigen 85b stim resembles ESAT-6 stimulation for IFN-γ and IL-17A staining of CD4^+^ T cells. As in Figure 3, WT B6 mice were aerosol infected with either WT, Δ*eccC1,* Δ*eccC1::eccC1*, *fadD28::tn*, or *fadD28::tn* + p*fadD28 M. tuberculosis* Erdman strain. 21 days post infection, mice were sacrificed, lung single cell homogenates were stimulated with Antigen 85b peptide pool and measured for lung **(A)** IFN-γ^+^ CD4^+^ T cells and **(B)** IL-17A^+^ CD4^+^ T cells by flow cytometry. Results in A and B are representative of 2 independent experiments. *, p < 0.05;**, p < 0.01 (unpaired nonparametric Mann-Whitney U test).

**Figure S5:**
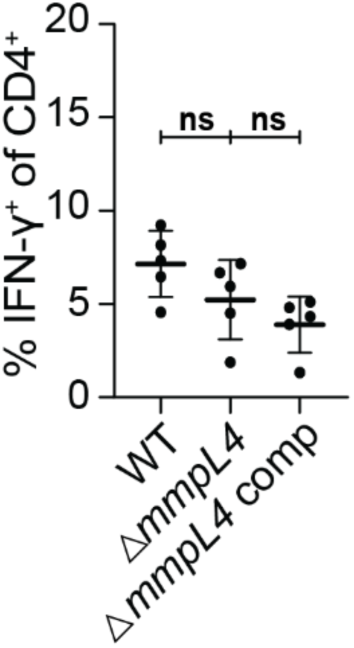
Δ*mmpL4* Mtb induces similar IFN-γ^+^ CD4^+^ T cells as WT or complemented Mtb. Mice were aerosol infected with WT, Δ*mmpL4* mutant, or complemented Δ*mmpL4* Mtb Erdman strain and evaluated for IFN-γ^+^ CD4^+^ T cells after restimulation with ESAT-6 peptide by flow cytometry. Results are representative of 2 independent experiments. *, p < 0.05; **, p < 0.01 (unpaired nonparametric Mann-Whitney U test).

**Figure S6:**
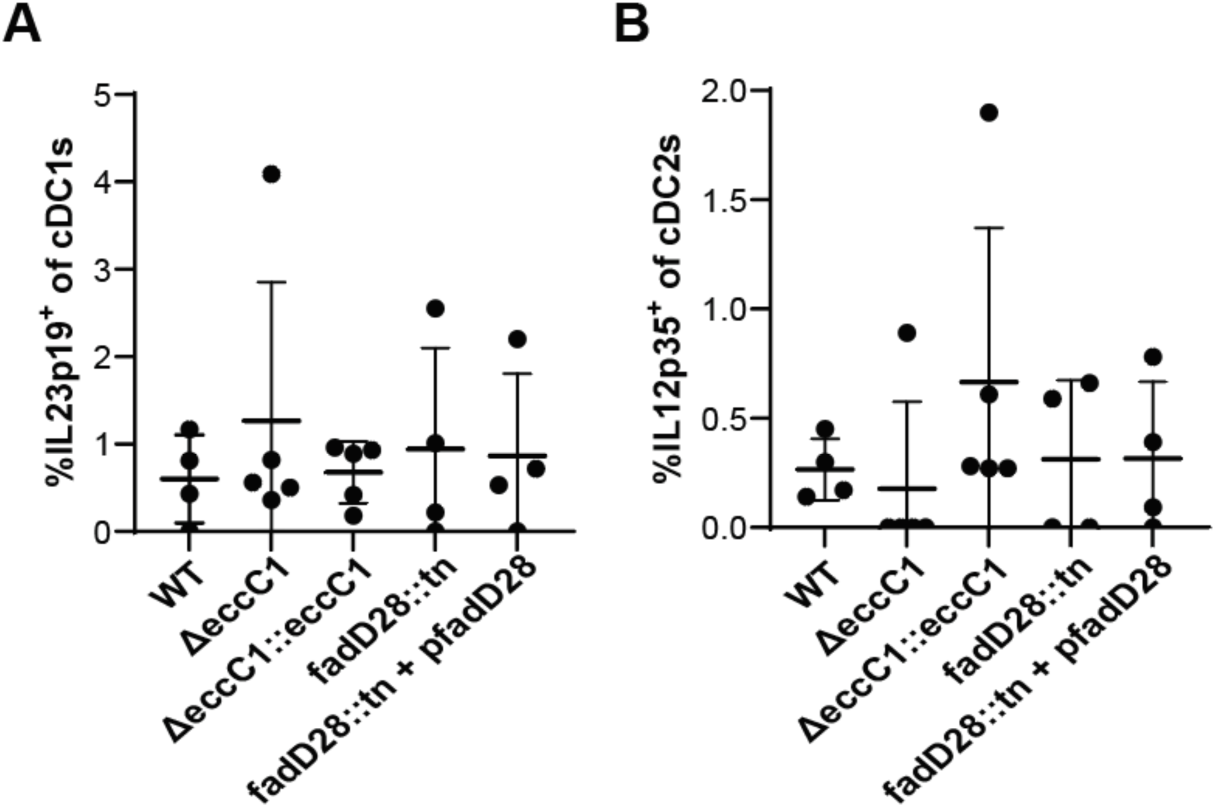
There is little to no IL-23 p19 expression in cDC1s and IL-12 p35 expression in cDC2s during Mtb infection. As in Figure 5, WT B6 mice were aerosol infected with WT, Δ*eccC1,* Δ*eccC1::eccC1*, *fadD28::tn*, or *fadD28::tn* + p*fadD28 M. tuberculosis* Erdman strain. 21 days post infection, mice were sacrificed, mediastinal lymph nodes were extracted and processed for ICS. Analysis of **(A)** IL-23 p19^+^ cDC1s and **(B)** IL-12 p35^+^ cDC2s by flow cytometry. Results in A and B are representative of 2 independent experiments. (unpaired nonparametric Mann-Whitney U test).

## References

1. World Health Organization, Global Tuberculosis Report 2022. *Geneva:* World Health Organization. 2022. Global Tuberculosis Report 2022. (2022). Available at: https://www.who.int/publications/i/item/9789240061729.

2. P. E. Fine, Variation in protection by BCG: implications of and for heterologous immunity. Lancet 346, 1339–1345 (1995).

3. P. Kumar, Corrigendum: A perspective on the success and failure of BCG. Front. Immunol. 13, 850325 (2022).

4. F. Zhou, D. Zhang, Recent advance in the development of tuberculosis vaccines in clinical trials and virus-like particle-based vaccine candidates. Front. Immunol. 14, 1238649 (2023).

5. J. L. Flynn, et al., An essential role for interferon gamma in resistance to Mycobacterium tuberculosis infection. J. Exp. Med. 178, 2249–2254 (1993).

6. J.-L. Casanova, L. Abel, Genetic dissection of immunity to mycobacteria: the human model. Annu. Rev. Immunol. 20, 581–620 (2002).

7. M. D. Tameris, et al., Safety and efficacy of MVA85A, a new tuberculosis vaccine, in infants previously vaccinated with BCG: a randomised, placebo-controlled phase 2b trial. Lancet 381, 1021–1028 (2013).

8. B. Zhang, et al., Association between IL-18, IFN-γ and TB susceptibility: a systematic review and meta-analysis. Ann. Palliat. Med. 10, 10878–10886 (2021).

9. L. Guglani, S. A. Khader, Th17 cytokines in mucosal immunity and inflammation. Curr. Opin. HIV AIDS 5, 120–127 (2010).

10. S. L. Gaffen, Structure and signalling in the IL-17 receptor family. Nat. Rev. Immunol. 9, 556–567 (2009).

11. V. Lazarevic, et al., T-bet represses T(H)17 differentiation by preventing Runx1-mediated activation of the gene encoding RORγt. Nat. Immunol. 12, 96–104 (2011).

12. M. Pepper, et al., Different routes of bacterial infection induce long-lived T. Nat. Immunol. 11, 83–89 (2009).

13. W.-I. Yeh, I. L. McWilliams, L. E. Harrington, IFNγ inhibits Th17 differentiation and function via Tbet-dependent and Tbet-independent mechanisms. J. Neuroimmunol. 267, 20–27 (2014).

14. B. Guo, E. Y. Chang, G. Cheng, The type I IFN induction pathway constrains Th17-mediated autoimmune inflammation in mice. J. Clin. Invest. 118, 1680–1690 (2008).

15. H. P. Gideon, et al., Multimodal profiling of lung granulomas in macaques reveals cellular correlates of tuberculosis control. Immunity 55, 827–846.e10 (2022).

16. P. Ogongo, et al., High-parameter phenotypic characterization reveals a subset of human Th17 cells that preferentially produce IL-17 against M. tuberculosis antigen. Front. Immunol. 15, 1378040 (2024).

17. M. Sun, et al., Specific CD4+ T cell phenotypes associate with bacterial control in people who “resist” infection with Mycobacterium tuberculosis. Nat. Immunol. 25, 1411–1421 (2024).

18. A. Nathan, et al., Multimodally profiling memory T cells from a tuberculosis cohort identifies cell state associations with demographics, environment and disease. Nat. Immunol. 22, 781–793 (2021).

19. R. Gopal, et al., Unexpected role for IL-17 in protective immunity against hypervirulent Mycobacterium tuberculosis HN878 infection. PLoS Pathog. 10, e1004099 (2014).

20. S. A. Khader, et al., IL-23 and IL-17 in the establishment of protective pulmonary CD4+ T cell responses after vaccination and during Mycobacterium tuberculosis challenge. Nat. Immunol. 8, 369–377 (2007).

21. N. Aguilo, et al., Pulmonary but Not Subcutaneous Delivery of BCG Vaccine Confers Protection to Tuberculosis-Susceptible Mice by an Interleukin 17-Dependent Mechanism. J. Infect. Dis. 213, 831–839 (2016).

22. R. M. Jong, et al., Mucosal Vaccination with Cyclic Dinucleotide Adjuvants Induces Effective T Cell Homing and IL-17-Dependent Protection against Mycobacterium tuberculosis Infection. J. Immunol. 208, 407–419 (2022).

23. S. A. Stanley, S. Raghavan, W. W. Hwang, J. S. Cox, Acute infection and macrophage subversion by Mycobacterium tuberculosis require a specialized secretion system. Proc. Natl. Acad. Sci. U. S. A. 100, 13001–13006 (2003).

24. S. A. Stanley, J. E. Johndrow, P. Manzanillo, J. S. Cox, The Type I IFN response to infection with Mycobacterium tuberculosis requires ESX-1-mediated secretion and contributes to pathogenesis. Mian Yi Xue Za Zhi 178, 3143–3152 (2007).

25. R. O. Watson, et al., The Cytosolic Sensor cGAS Detects Mycobacterium tuberculosis DNA to Induce Type I Interferons and Activate Autophagy. Cell Host Microbe 17, 811–819 (2015).

26. R. O. Watson, P. S. Manzanillo, J. S. Cox, Extracellular M. tuberculosis DNA targets bacteria for autophagy by activating the host DNA-sensing pathway. Cell 150, 803–815 (2012).

27. J. S. Cox, B. Chen, M. McNeil, W. R. Jacobs, Complex lipid determines tissue-specific replication of Mycobacterium tuberculosis in mice. Nature 402, 79–83 (1999).

28. A. K. Barczak, et al., Systematic, multiparametric analysis of Mycobacterium tuberculosis intracellular infection offers insight into coordinated virulence. PLoS Pathog. 13, e1006363 (2017).

29. A. Schlitzer, et al., IRF4 transcription factor-dependent CD11b+ dendritic cells in human and mouse control mucosal IL-17 cytokine responses. Immunity 38, 970–983 (2013).

30. P. Soldevilla, C. Vilaplana, P.-J. Cardona, Mouse models for Mycobacterium tuberculosis pathogenesis: Show and do not tell. Pathogens 12, 49 (2022).

31. P. Andersen, A. B. Andersen, A. L. Sørensen, S. Nagai, Recall of long-lived immunity to Mycobacterium tuberculosis infection in mice. J. Immunol. 154, 3359–3372 (1995).

32. A. Bekmurzayeva, M. Sypabekova, D. Kanayeva, Tuberculosis diagnosis using immunodominant, secreted antigens of Mycobacterium tuberculosis. Tuberculosis (Edinb.) 93, 381–388 (2013).

33. M. Karbalaei Zadeh Babaki, S. Soleimanpour, S. A. Rezaee, Antigen 85 complex as a powerful Mycobacterium tuberculosis immunogene: Biology, immune-pathogenicity, applications in diagnosis, and vaccine design. Microb. Pathog. 112, 20–29 (2017).

34. T. Dileepan, et al., Robust antigen specific th17 T cell response to group A Streptococcus is dependent on IL-6 and intranasal route of infection. PLoS Pathog. 7, e1002252 (2011).

35. D. Sleep, J. Cameron, L. R. Evans, Albumin as a versatile platform for drug half-life extension. Biochim. Biophys. Acta 1830, 5526–5534 (2013).

36. B. M. Sullivan, et al., Increased susceptibility of mice lacking T-bet to infection with Mycobacterium tuberculosis correlates with increased IL-10 and decreased IFN-gamma production. Mian Yi Xue Za Zhi 175, 4593–4602 (2005).

37. M. A. Sallin, et al., Th1 Differentiation Drives the Accumulation of Intravascular, Non-protective CD4 T Cells during Tuberculosis. Cell Rep. 18, 3091–3104 (2017).

38. A. E. Hinman, et al., Mycobacterium tuberculosis canonical virulence factors interfere with a late component of the TLR2 response. Elife 10 (2021).

39. J. Augenstreich, et al., ESX-1 and phthiocerol dimycocerosates of Mycobacterium tuberculosis act in concert to cause phagosomal rupture and host cell apoptosis. Cell. Microbiol. 19 (2017).

40. T. D. Crosskey, K. S. H. Beckham, M. Wilmanns, The ATPases of the mycobacterial type VII secretion system: Structural and mechanistic insights into secretion. Prog. Biophys. Mol. Biol. 152, 25–34 (2020).

41. R. M. Wells, et al., Discovery of a siderophore export system essential for virulence of Mycobacterium tuberculosis. PLoS Pathog. 9, e1003120 (2013).

42. V. S. Ramgolam, Y. Sha, J. Jin, X. Zhang, S. Markovic-Plese, IFN-beta inhibits human Th17 cell differentiation. Mian Yi Xue Za Zhi 183, 5418–5427 (2009).

43. D. X. Ji, et al., Role of the transcriptional regulator SP140 in resistance to bacterial infections via repression of type I interferons. Elife 10 (2021).

44. C. Dye, P. Glaziou, K. Floyd, M. Raviglione, Prospects for tuberculosis elimination. Annu. Rev. Public Health 34, 271–286 (2013).

45. R. Madan-Lala, K. V. Peixoto, F. Re, J. Rengarajan, Mycobacterium tuberculosis Hip1 dampens macrophage proinflammatory responses by limiting toll-like receptor 2 activation. Infect. Immun. 79, 4828–4838 (2011).

46. J. K. Sia, E. Bizzell, R. Madan-Lala, J. Rengarajan, Engaging the CD40-CD40L pathway augments T-helper cell responses and improves control of Mycobacterium tuberculosis infection. PLoS Pathog. 13, e1006530 (2017).

47. A. B. Enriquez, et al., Mycobacterium tuberculosis impedes CD40-dependent notch signaling to restrict Th17 polarization during infection. iScience 25, 104305 (2022).

48. S. A. Khader, et al., IL-23 compensates for the absence of IL-12p70 and is essential for the IL-17 response during tuberculosis but is dispensable for protection and antigen-specific IFN-gamma responses if IL-12p70 is available. J. Immunol. 175, 788–795 (2005).

49. S. Sakai, et al., CD4 T Cell-Derived IFN-γ Plays a Minimal Role in Control of Pulmonary Mycobacterium tuberculosis Infection and Must Be Actively Repressed by PD-1 to Prevent Lethal Disease. PLoS Pathog. 12, e1005667 (2016).

50. I. Comas, et al., Human T cell epitopes of Mycobacterium tuberculosis are evolutionarily hyperconserved. Nat. Genet. 42, 498–503 (2010).

51. P. Ogongo, et al., Rare Variable M. tuberculosis Antigens induce predominant Th17 responses in human infection. bioRxivorg (2024).

52. F. McNab, K. Mayer-Barber, A. Sher, A. Wack, A. O’Garra, Type I interferons in infectious disease. Nat. Rev. Immunol. 15, 87–103 (2015).

53. S. L. Gaffen, R. Jain, A. V. Garg, D. J. Cua, The IL-23-IL-17 immune axis: from mechanisms to therapeutic testing. Nat. Rev. Immunol. 14, 585–600 (2014).

54. G. L. Stritesky, N. Yeh, M. H. Kaplan, IL-23 promotes maintenance but not commitment to the Th17 lineage. J. Immunol. 181, 5948–5955 (2008).

55. J. G. Krueger, et al., IL-23 past, present, and future: a roadmap to advancing IL-23 science and therapy. Front. Immunol. 15, 1331217 (2024).

56. L. Solans, et al., A specific polymorphism in Mycobacterium tuberculosis H37Rv causes differential ESAT-6 expression and identifies WhiB6 as a novel ESX-1 component. Infect. Immun. 82, 3446–3456 (2014).

57. R. Stadhouders, E. Lubberts, R. W. Hendriks, A cellular and molecular view of T helper 17 cell plasticity in autoimmunity. J. Autoimmun. 87, 1–15 (2018).

58. G. G. Mahairas, P. J. Sabo, M. J. Hickey, D. C. Singh, C. K. Stover, Molecular analysis of genetic differences between Mycobacterium bovis BCG and virulent M. bovis. J. Bacteriol. 178, 1274–1282 (1996).

59. T. Hsu, et al., The primary mechanism of attenuation of bacillus Calmette-Guerin is a loss of secreted lytic function required for invasion of lung interstitial tissue. Proc. Natl. Acad. Sci. U. S. A. 100, 12420–12425 (2003).

60. A. Arbues, et al., Construction, characterization and preclinical evaluation of MTBVAC, the first live-attenuated M. tuberculosis-based vaccine to enter clinical trials. Vaccine 31, 4867–4873 (2013).

61. K. Dijkman, et al., Pulmonary MTBVAC vaccination induces immune signatures previously correlated with prevention of tuberculosis infection. Cell Rep. Med. 2, 100187 (2021).

